# Essential roles of the ANKRD31-REC114 interaction in meiotic recombination and mouse spermatogenesis

**DOI:** 10.1101/2023.04.27.538541

**Authors:** Jiaqi Xu, Tao Li, Soonjoung Kim, Michiel Boekhout, Scott Keeney

## Abstract

Meiotic DNA double-strand breaks (DSBs) initiate homologous recombination and are crucial for ensuring proper chromosome segregation. In mice, ANKRD31 recently emerged as a regulator of DSB timing, number, and location, with a particularly important role in targeting DSBs to the pseudoautosomal regions (PARs) of sex chromosomes. ANKRD31 interacts with multiple proteins, including the conserved and essential DSB-promoting factor REC114, so it was hypothesized to be a modular scaffold that “anchors” other proteins together and to meiotic chromosomes. To determine if and why the REC114 interaction is important for ANKRD31 function, we generated mice with *Ankrd31* mutations that either reduced (missense mutation) or eliminated (C-terminal truncation) the ANKRD31– REC114 interaction without diminishing contacts with other known partners. A complete lack of the ANKRD31–REC114 interaction mimicked an *Ankrd31* null, with delayed DSB formation and recombination, defects in DSB repair, and altered DSB locations including failure to target DSBs to the PARs. In contrast, when the ANKRD31– REC114 interaction was substantially but not completely disrupted, spermatocytes again showed delayed DSB formation globally, but recombination and repair were hardly affected and DSB locations were similar to control mice. The missense *Ankrd31* allele showed a dosage effect, wherein combining it with the null or C-terminal truncation allele resulted in intermediate phenotypes for DSB formation, recombination, and DSB locations. Our results show that ANKRD31 function is critically dependent on its interaction with REC114, and that defects in ANKRD31 activity correlate with the severity of the disruption of the interaction.

**Significance:** Homologous recombination initiated by double-strand breaks (DSBs) during meiosis is a nearly universal feature of eukaryotic lifecycles, but is also dangerous because DSBs are potentially toxic or mutagenic. The vertebrate-specific protein ANKRD31 is an important regulator of DSB formation, proposed to be a scaffold protein that coordinates the activities of multiple DSB-promoting factors, including the widely conserved REC114. We test this hypothesis here through generation of targeted *Ankrd31* mutations that specifically attenuate or eliminate the ANKRD31-REC114 interaction. Analysis of this allelic series demonstrates that the ANKRD31-REC114 interaction is essential for all ANKRD31 activities in vivo, providing insight into how ANKRD31 controls DSB locations, timing, and number.

## Introduction

During meiosis, one round of DNA replication is followed by two rounds of chromosome segregation to reduce the chromosome complement. In many species, the segregation of homologous chromosomes during meiosis I requires programmed DNA double-strand breaks (DSBs) to be generated and then repaired by homologous recombination. These DSBs are formed by SPO11 and accessory proteins (1), which in mice include meiosis-specific REC114, MEI4, and IHO1 (orthologs of yeast Rec114, Mei4, and Mer2, respectively) (2-5). Although the major protein players have been known for some time, the physical and functional interactions among them are not well understood, particularly in mammals.

The vertebrate-specific protein ANKRD31 (Ankyrin Repeat Domain Containing 31) was recently identified as a direct interaction partner of REC114 (6, 7). ANKRD31 contains two ankyrin repeat domains and three additional conserved regions (**Figure 1A**). The C-terminal-most conserved region (aa 1808–1857) wraps extensively around the N-terminal pleckstrin homology (PH) domain of REC114 as shown by a crystal structure of the complex (6) (**Figure S1A**). Other segments in ANKRD31 interact with additional partners IHO1, MEI1, PTIP, and ZMYM3 (6-8) (**Figure 1A**,**B**).

**Figure 1.**
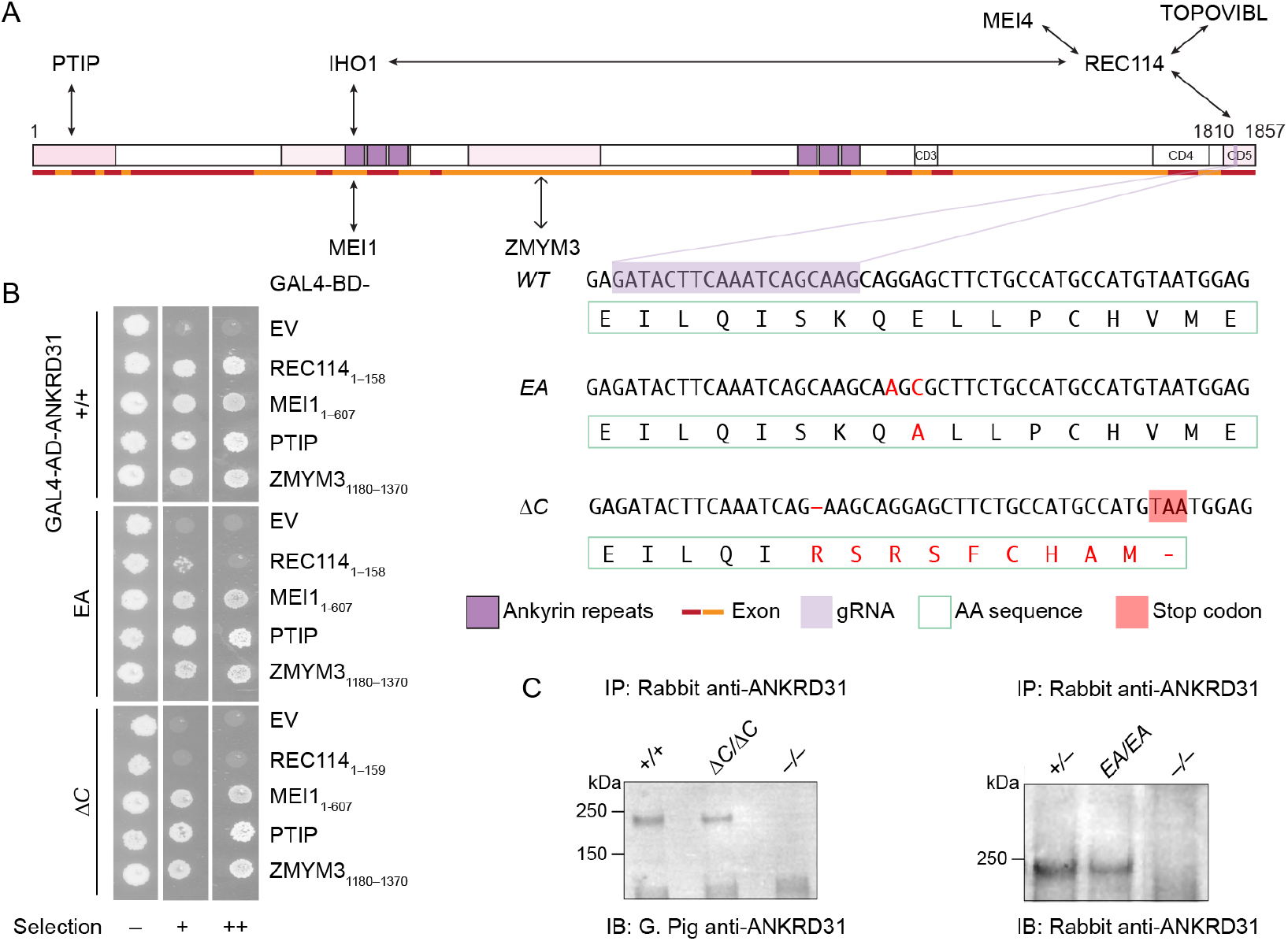
Generation of ANKRD31–REC114 interaction-deficient mice. (A) Diagram of mouse ANKRD31 domain structure, interactions and CRISPR/Cas9 guide RNA target. Ankyrin repeat domains (dark purple), conserved domains (CD3–CD5), interacting regions (shaded in pink), exons (red and orange lines), and gRNA sequence (light purple) are indicated. DNA and amino acid sequences of wild-type and mutated alleles are indicated. Mutant sequences are shown in red. The premature stop codon in ANKRD31-ΔC is shaded in red. (B) Y2H interactions of wild-type and mutated ANKRD31 with full-length or parts (indicated) of REC114, MEI1, PTIP, and ZMYM3. Cells express the indicated Gal4 activating domain (AD) and binding domain (BD) fusions. EV, empty vector. “Selection” indicates amino acid dropouts and aureobasidin to detect reporter activation at moderate (+) and high (++) stringency. Immunoprecipitation (IP) and immunoblotting (IB) of ANKRD31 from whole-testis extracts of control and mutated mice. Adult mice (> 2 mos old) were used in the experiment on the right. Younger mutant mice (1.5 mos old) were used for the experiment on the left to reduce potential effects of the altered testis cellularity in *Ankrd31*^*ΔC/ΔC*^ adults (Methods).

*Ankrd31*-deficient male mice have delayed DSB formation and defects in DSB repair, accompanied by partially penetrant arrest and apoptosis of spermatocytes during the pachytene stage (6, 7). Because oocytes also show these DSB defects, *Ankrd31*^*−/−*^ females are fertile when young, but have reduced oocyte reserve and show premature ovarian failure (6, 7).

*Ankrd31* deficiency also leads to two major changes in DSB locations. First, there is a near complete loss of the high-frequency DSBs in the pseudoautosomal regions (PARs) of the X and Y chromosomes, which are necessary for sex chromosome pairing, recombination, and segregation in spermatocytes (6-8). Consequently, most cells that successfully develop beyond pachynema arrest at metaphase I because of achiasmate sex chromosomes, leading to sterility in males. Second, there is a change in global DSB locations (6, 7), in which the DSB hotspots that are targeted by the PRDM9 histone methyltransferase (9) are joined by additional (“default”) hotspots that occur near promoters and other genomic locations that are typically only targeted for SPO11 activity in the absence of PRDM9 function (10).

During early prophase I, ANKRD31 forms numerous small foci that colocalize with DSB-promoting factors such as REC114, MEI1, and MEI4 on chromosome axes (6-8). These small foci are presumably involved in forming the hundreds of DSBs that are targeted by PRDM9 across the genome. Alterations at these focal sites are also presumably the cause of the changed DSB locations, delayed DSB formation, and defective DSB repair in *Ankrd31* null mutants. In addition, ANKRD31 forms large immunostaining “blobs” on repetitive arrays of the mo-2 minisatellite that are present on the PAR and the centromere-distal ends of chromosomes 4, 9 and 13 (6-8). These blobs also contain REC114, MEI4, MEI1 and IHO1, and are responsible for PAR axis remodeling that enables efficient DSB formation (8). ANKRD31, REC114 and MEI4—but not IHO1—are essential for mo-2 blob formation (6-8).

Given the complexity of ANKRD31 function and the fact that ANKRD31 interacts with multiple meiotic proteins, we hypothesized that ANKRD31 might act as a scaffold that anchors REC114 and other interactors to specific genomic loci at different times, thus regulating DSB formation (6). A key prediction of this hypothesis is that each specific ANKRD31 protein-protein interaction is important for meiosis, but which function(s) of ANKRD31 depends on which interactions is unknown.

To address these issues, we used CRISPR/Cas9 genome editing to alter the last *Ankrd31* exon (encoding the REC114-interaction domain) to generate mutations that disrupt the ANKRD31–REC114 interaction to different extents. Analysis of mice carrying *Ankrd31* mutant alleles in various combinations showed that this interaction is essential for all ANKRD31 functions, with specific functions dependent to different quantitative degrees on the strength of the interaction.

## Results

### Generation of ANKRD31–REC114 interaction-deficient mice

In the crystal structure for ANKRD31 residues 1808–1857 (ANKRD31_C_) complexed with REC114 residues 1–158 (REC114_N_), the side chain of ANKRD31 Glu-1831 is anchored by two intermolecular salt bridges and one hydrogen bond (6) (**Figure S1A**). In pulldown assays with recombinant GST-tagged ANKRD31_C_ and purified REC114_N_, substituting alanine for Glu-1831 dramatically reduced the interaction (6). This mutation similarly disrupted the yeast two-hybrid (Y2H) interaction between ANKRD31_C_ and REC114 1–145 (**Figure S1B**).

We therefore used a guide RNA (**Figure 1A**) to target Glu-1831 encoded in exon 25 plus a donor DNA containing desired mutations as the repair template to generate a point mutated allele harboring G-to-A and A-to-C mutations at positions 5490 and 5492 (an E to A mutation at residue 1831, hereafter *Ankrd31*^*EA*^). We also fortuitously obtained a truncated allele (hereafter *Ankrd31*^*ΔC*^) with a 1-bp deletion at position 5484 that results in a frameshift mutation and premature termination, replacing the last 21 amino acids with 9 extraneous residues (**Figure 1A**).

We performed Y2H assays to test if the interaction with REC114_N_ is disrupted by these two mutations when in the context of full-length ANKRD31 protein. EA severely, but not completely, disrupted the interaction, while ANKRD31-ΔC showed no evidence of residual interaction (**Figure 1B**). The less complete interaction defect for the point mutation when it was in the full-length protein as compared to the short ANKRD31_C_ peptide (6) (**Figure S1B**) suggests that there may be additional contacts between REC114_N_ and other parts of ANKRD31 (7). If so, the inability of otherwise full-length ANKRD31-ΔC to interact with REC114_N_ indicates that these contacts are not sufficient.

Both ANKRD31-EA and ANKRD31-ΔC proteins were detected at normal levels by immunoprecipitation from whole-testis extracts followed by immunoblotting (**Figure 1C**), indicating that the mutations did not substantially alter protein stability in vivo. In addition, both mutant proteins maintained Y2H interaction with other known ANKRD31 partners MEI1, PTIP, and ZMYM3 (**Figure 1B**), which interact with other parts of ANKRD31 (**Figure 1A**) (8). We conclude that we successfully generated mutations that cause defects specifically in the ANKRD31– REC114 interaction to different extents.

### Hypogonadism and infertility in an interaction-deficient Ankrd31 allelic series

*Ankrd31*^*–/–*^ males are sterile and their testes are about one-third of the weight of those in control mice (6, 7) (**Figure 2A**). *Ankrd31*^*ΔC/ΔC*^ males were also sterile: neither of the two animals tested (9 mos old) sired offspring when bred with fertile females for 8 weeks, with similarly sized testes as in *Ankrd31*^*–/–*^ males (**Figure 2A**). In contrast, *Ankrd31*^*EA/EA*^ males were fertile, albeit with testis weights significantly lower than wild type (p < 0.0001, Student’s t test; **Figure 2A**). Consistent with fertility, epididymal sperm were observed in *Ankrd31*^*EA/EA*^ and control mice, but not in *Ankrd31*^*ΔC/ΔC*^ or *Ankrd31*^*–/–*^ males (**Figure 2B**).

**Figure 2.**
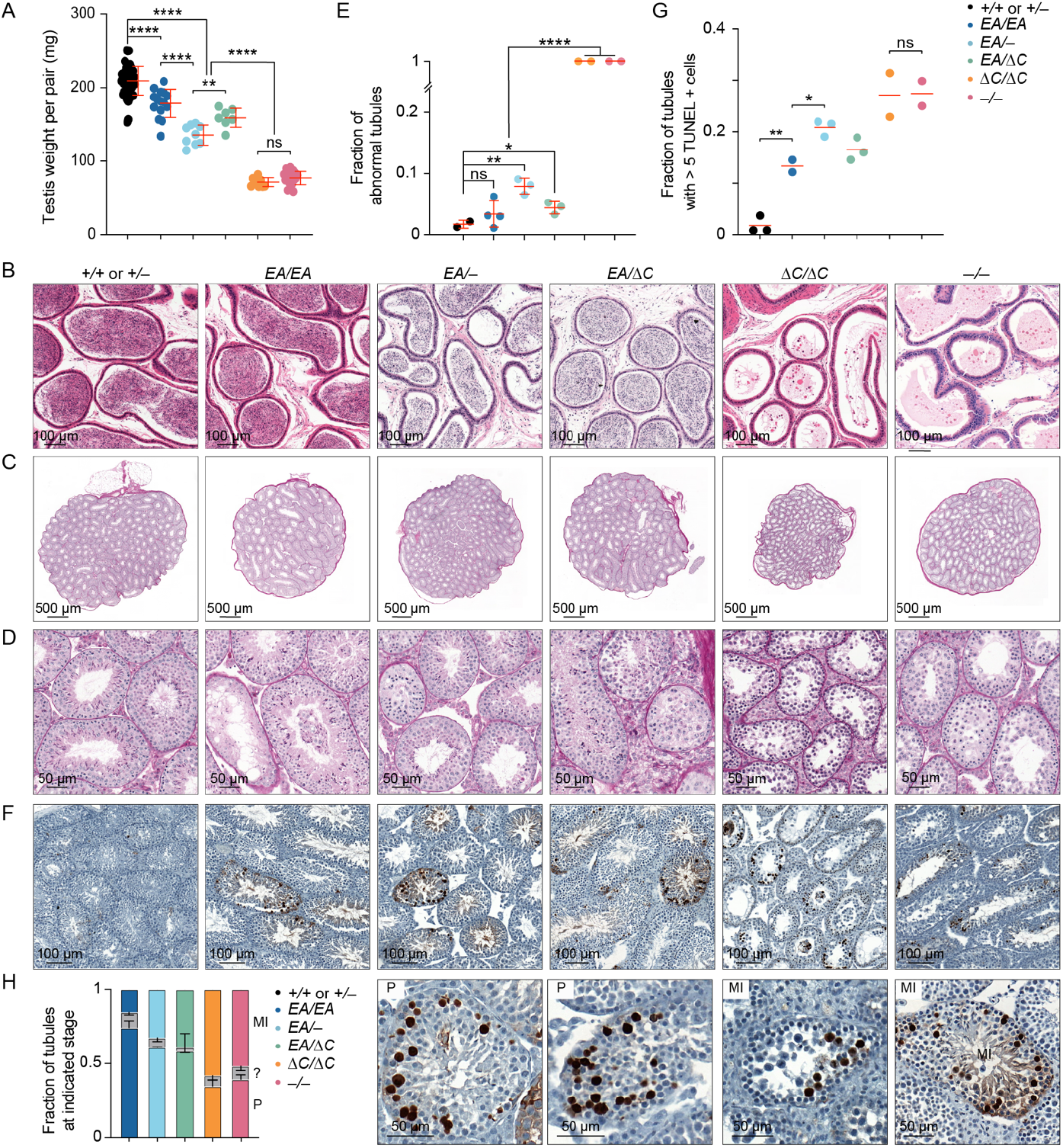
Hypogonadism and infertility in an interaction-deficient *Ankrd31* allelic series. (A) Quantification of testis weights (red lines, means ± SD). Mice were 2–11 mos old. (B) Sections of epididymides from adult mice (2–8 mos old), PFA fixed and H&E stained. (C,D) Sections of adult (2–8 mos old) testes (C) and seminiferous tubules (D), Bouin’s fixed and periodic acid Schiff (PAS) stained. (E) Quantification of tubules with abnormal spermatogenesis. Each point on the scatterplot is the measurement from one animal; the red lines are means. (F,G) Adult (2–8 mos old) testis sections were stained with TUNEL and hematoxylin. Each point on the scatterplot is the measurement from one animal; the red lines are means. (H) Fraction of apoptotic tubules in mutants according to spermatocyte stage present (mean and SD for at least two experiments). MI, metaphase I; P, pachytene; ?, ambiguous. Representative examples of tubules with pachytene or metaphase I apoptosis are shown at right. The results of two-tailed Mann-Whitney U tests in A, E, G are shown: ns, not significant (p > 0.05), ∗p ≤ 0.05, ∗∗p ≤ 0.01, ∗∗∗p ≤ 0.001, and ∗∗∗∗p ≤ 0.0001. Underlying data for all plots in this and subsequent figures, including exact p values, are provided in **Supplemental File 1**.

To explore if there is a dosage effect of the *Ankrd31*^*EA*^ allele, we also generated compound heterozygous mutants *Ankrd31*^*EA/–*^ and *Ankrd31*^*EA/ΔC*^. Heterozygous males of both genotypes had testis weights that were significantly lower than in *Ankrd31*^*EA/EA*^ mice (p < 0.0001 and p = 0.0244, respectively), but significantly heavier than *Ankrd31*^*ΔC/ΔC*^ and *Ankrd31*^*−/−*^ males (p < 0.0001; **Figure 2A**). Epididymal sperm were observed in *Ankrd31*^*EA/–*^ and *Ankrd31*^*EA/ΔC*^ males (**Figure 2B**), and males of both genotypes were able to sire pups, although we cannot rule out that they were sub-fertile.

Testis sections from control animals had the full array of spermatogenic cells, including spermatocytes and round and elongated spermatids, as expected (**Figure 2C–D**). In contrast, *Ankrd31*^*−/−*^ tubules, as was previously shown (6, 7), contained spermatogonia and primary spermatocytes but were largely if not completely devoid of post-meiotic cells. *Ankrd31*^*ΔC/ΔC*^ tubules were again similar to *Ankrd31*^*−/−*^ (**Figure 2C–E**). In contrast, the majority of *Ankrd31*^*EA/EA*^ tubules appeared normal and only a small fraction appeared abnormal (emptier and lacking post-meiotic cells; **Figure 2C–E**). Compound heterozygotes *Ankrd31*^*EA/–*^ and *Ankrd31*^*EA/ΔC*^ both had some normal tubules but also had a larger fraction of abnormal tubules as compared to *Ankrd31*^*EA/EA*^ (**Figure 2C–E**).

Meiotic recombination defects cause hypogonadism and sterility because of spermatocyte apoptosis (11). We interpret the less populated tubules in interaction-deficient mutants as those in which apoptosis has already eliminated aberrant cells. Pachytene arrest can be triggered by persistent DSBs or defects in synapsis or meiotic sex chromosome inactivation (12-14). In contrast, metaphase I arrest is typical for mutants that can complete DSB repair but harbor achiasmate chromosomes that trigger a spindle checkpoint (15, 16).

As previously reported, TUNEL staining detected a high frequency (nearly 30%) of apoptotic tubules in *Ankrd31*^*−/−*^ testis sections, displaying partially penetrant pachytene arrest and more fully penetrant metaphase I arrest (6, 7) (**Figure 2F–G**). *Ankrd31*^*ΔC/ΔC*^ testis sections were comparable (**Figure 2F–G**). In contrast, *Ankrd31*^*EA/EA*^ testes had a smaller fraction (∼10%) of apoptotic tubules, and the compound heterozygotes again showed an intermediate phenotype (∼20% apoptotic tubules) (**Figure 2F–G**). In all mutants tested, dying spermatocytes were observed in both pachynema and metaphase I, indicating varying penetrance of arrest at both stages of prophase I (**Figure 2H**).

### ANKRD31–REC114 interaction deficiencies cause graded defects in DSB formation and recombination

To assess meiotic DSB formation and recombination, we immunostained chromosomes for γH2AX, RPA2, DMC1, and RAD51. γH2AX is a phosphorylated form of histone H2AX that is generated by the kinases ATM and ATR during meiotic prophase I in response to SPO11-generated DSBs (17-20). γH2AX initially appears genome-wide during early prophase I, then concentrates in a DSB-independent manner on the silenced X and Y chromosomes (18, 20, 21). RPA2 is a subunit of the single-stranded DNA (ssDNA) binding protein RPA (22). DMC1 (meiosis-specific) and RAD51 (ubiquitously expressed) are strand-exchange proteins homologous to bacterial RecA (22). The 3’-terminal ssDNA overhangs produced from meiotic DSBs are initially bound by RPA, which is then replaced by RAD51 and DMC1 (23-25). In normal meiosis, chromosome-associated foci of DMC1, RAD51 and RPA appear in leptonema, accumulate to maximal levels in early zygonema, then decline as DSB repair proceeds (26).

As previously shown, recombination focus numbers and γH2AX staining intensity were initially low in *Ankrd31*^*−/−*^ spermatocytes, but they eventually caught up to near normal levels (6, 7) (**Figures 3, 4A–B, and S2**). Seeing fewer RPA2 foci in early cells suggests that the reduction of RAD51 and DMC1 foci is caused by a delay and/or reduced efficiency of forming cytologically observable sites containing resected DSBs, not simply because of defective loading of the strand-exchange proteins. Thus, it has been interpreted that there is a delay in global DSB formation in the absence of ANKRD31 (6, 7). Additionally, elevated focus numbers and γH2AX staining were observed during pachynema (**Figures 3 and 4C–D**); these are interpreted as signs of persistent DSBs, suggesting a defect in DSB repair (6, 7).

**Figure 3.**
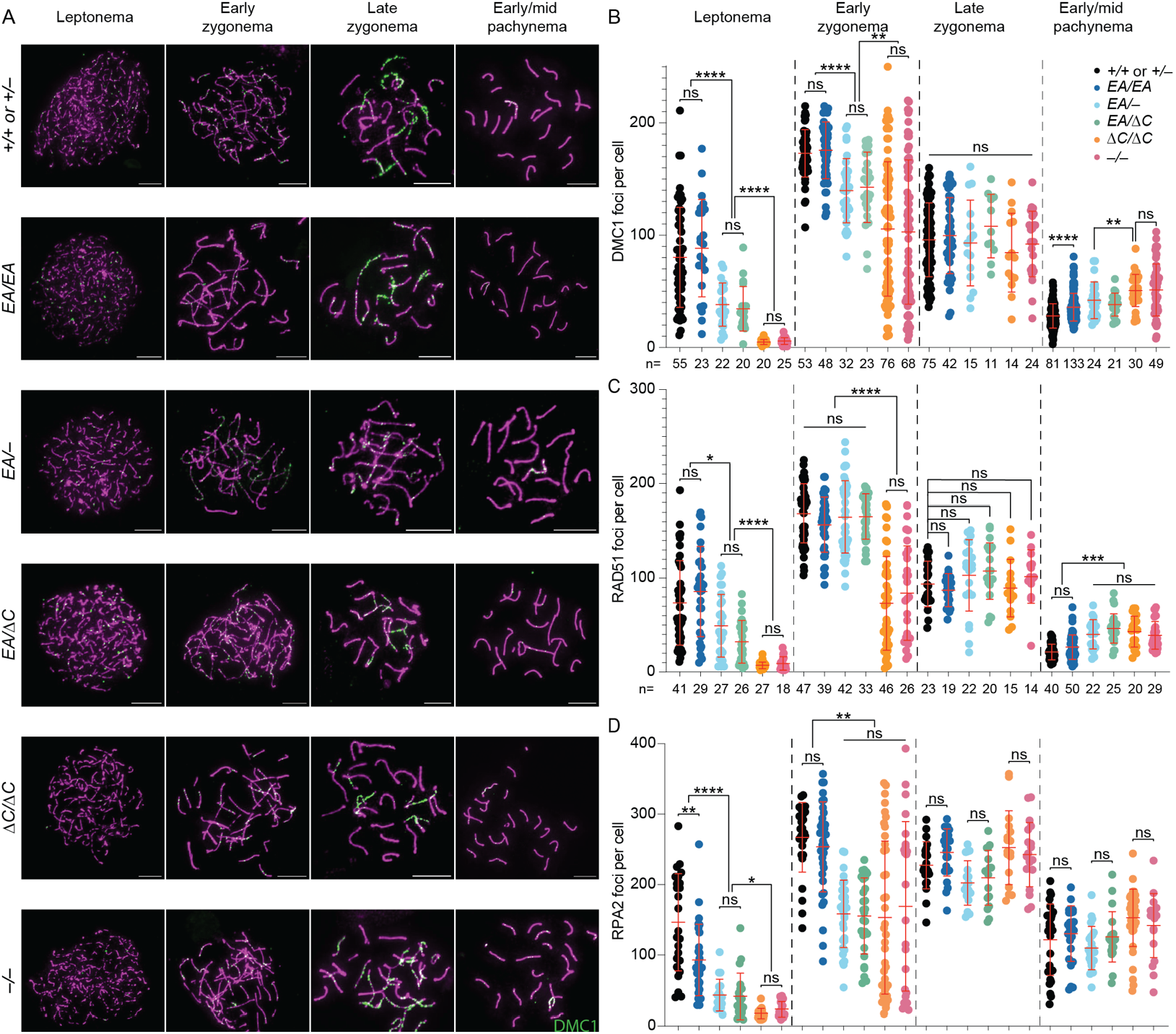
ANKRD31–REC114 interaction deficiencies cause progressive defects in DSB formation and recombination. (A) Representative DMC1 staining of spermatocyte chromosome spreads. Scale bars, 10 μm. (B–D) Quantification of focus numbers of DMC1 (B), RAD51 (C), and RPA2 (D). Each point is the count from one cell (total cell numbers are given below the graphs) from three or more animals of each genotype. The red lines are means ± SD. The results of two-tailed Mann-Whitney U tests are shown: ns, not significant (p > 0.05), ∗p ≤ 0.05, ∗∗p ≤ 0.01, ∗∗∗p ≤ 0.001, and ∗∗∗∗p ≤ 0.0001.

**Figure 4.**
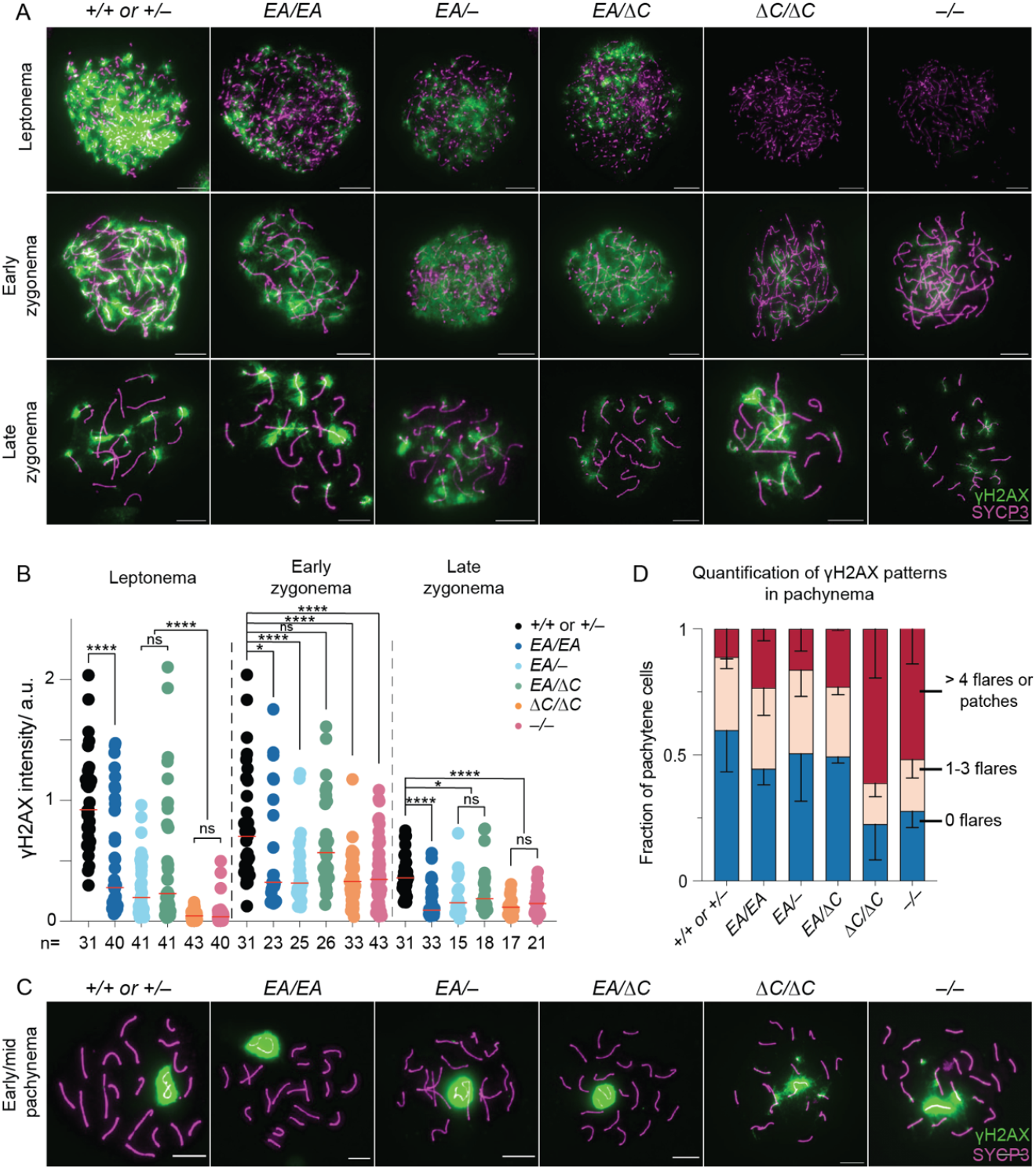
ANKRD31–REC114 interaction deficiencies cause progressive defects in DSB formation. (A) Representative γH2AX staining of spermatocyte chromosome spreads. Scale bars, 10 μm. (B) Quantification of γH2AX staining. The red lines are means. The results of two-tailed Mann-Whitney U tests are shown: ns, not significant (p > 0.05), ∗p ≤ 0.05, ∗∗p ≤ 0.01, ∗∗∗p ≤ 0.001, and ∗∗∗∗p ≤ 0.0001. (C) Representative γH2AX staining of spermatocyte chromosome spreads in pachynema. Scale bars, 10 μm. (D) Quantification of γH2AX patterns from at least two mice per genotype. Error bars indicate SD.

*Ankrd31*^*ΔC/ΔC*^ spermatocytes behaved comparably to *Ankrd31*^*−/−*^ spermatocytes for all of these DSB markers, at all substages of prophase I (**Figures 3, 4, and S2**). We conclude that the ANKRD31–REC114 interaction is essential for ANKRD31 functions in global DSB formation and recombination.

*Ankrd31*^*EA/EA*^ mice showed a similar but quantitatively milder defect. Compared to control mice, *Ankrd31*^*EA/EA*^ spermatocytes had reduced γH2AX staining throughout early prophase I (**Figure 4A–B**). RPA2 focus numbers were also lower during leptonema but caught up during later substages, while RAD51 and DMC1 foci remained comparable to controls throughout (**Figure 3 and S2**). Again, *Ankrd31*^*EA/–*^ and *Ankrd31*^*EA/ΔC*^ compound heterozygotes showed an intermediate phenotype, with reduced γH2AX staining throughout early prophase I (**Figure 4A–B**) and even fewer DMC1, RAD51, and RPA foci than the *Ankrd31*^*EA/EA*^ homozygous mice (**Figures 3 and S2**).

In all cases, focus numbers declined as chromosomes synapsed, but to different extents, resulting in more or less similar numbers of foci at late zygonema for all of the genotypes that include the missense allele. However, pachytene cells with apparently normal synapsis also had elevated numbers of DMC1 and RAD51 foci, with the degree of elevation correlating with the severity of the REC114 interaction defect (**Figure 3B–C**). We conclude that attenuating the ANKRD31–REC114 interaction causes a delay in global DSB formation and a defect in completion of recombination that are both quantitatively more modest than when the interaction is more completely disrupted.

We additionally examined crossing over on autosomes by staining for MLH1, a component of the Holliday junction resolvase (27, 28). Pachytene cells with complete autosome synapsis had similar numbers of MLH1 foci in all genotypes (means of 24.6 in wild type, 24.3 in *Ankrd31*^*EA/EA*^, 25.1 in *Ankrd31*^*EA/–*^, 24.3 in *Ankrd31*^*EA/ΔC*^, and 24.4 in *Ankrd31*^*ΔC/ΔC*^, and 24.8 in *Ankrd31*^*−/−*^; **Figure 5A–B**). This analysis did not confirm the small but statistically significant increase (6) or decrease (7) in MLH1 foci previously reported for *Ankrd31*^*−/−*^, but similar to previous results, a small increase in the number of cells with at least one autosome lacking an MLH1 focus was seen in all mutants (**Figure 5C**). The increases in *Ankrd31*^*ΔC/ΔC*^ and in *Ankrd31*^*−/−*^ were significant (p = 0.034 and 0.042, respectively; Fisher’s exact test).

**Figure 5.**
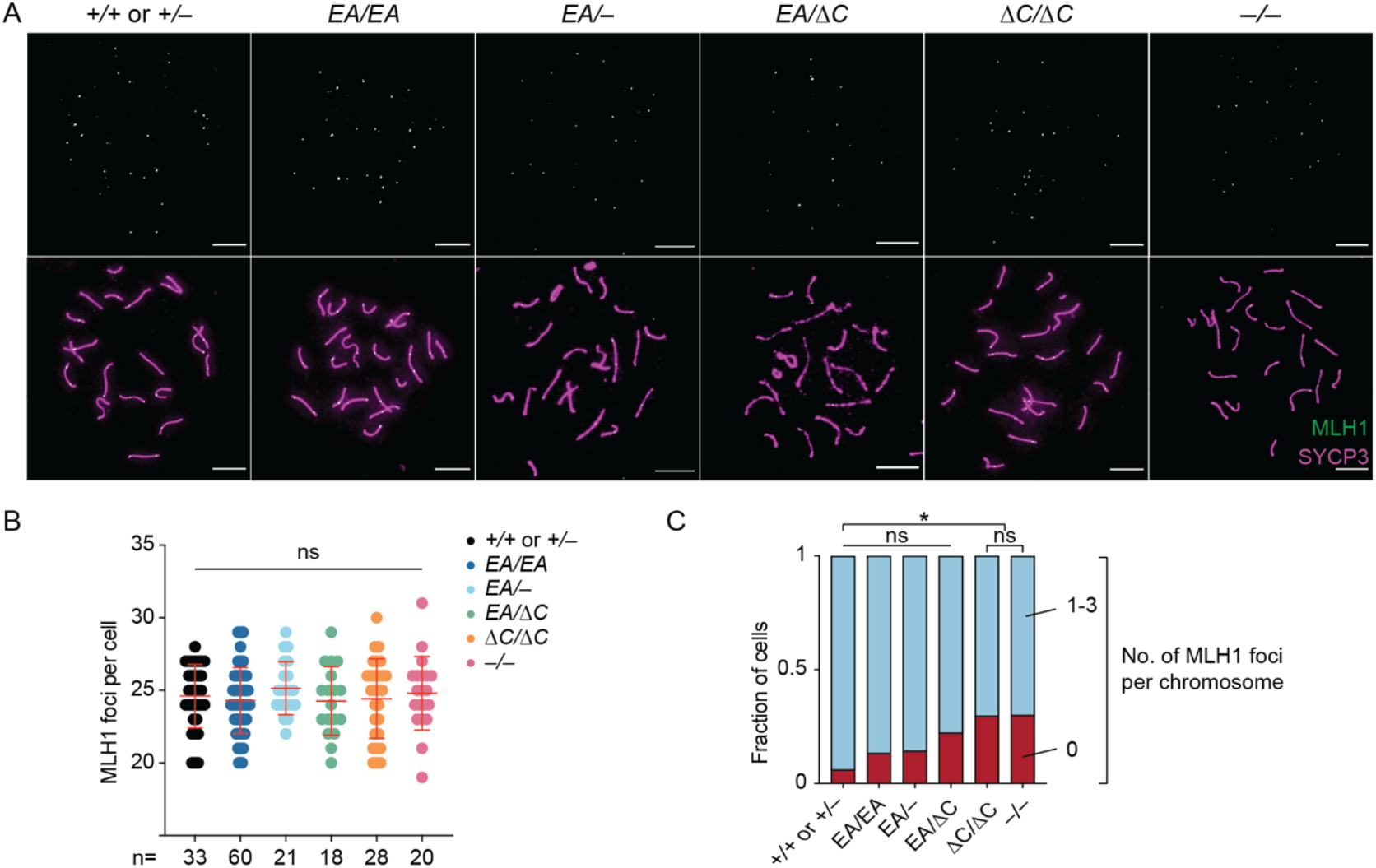
Autosomal recombination. (A) Representative pachytene cells labeled with anti-SYCP3 and anti-MLH1 antibodies. Scale bars, 10 μm. (B) Quantification of MLH1 foci from two or more animals of each genotype. The number of cells analyzed is indicated below. The red lines are means ± SD. The results of two-tailed Mann-Whitney U tests are shown: ns, not significant (p > 0.05). (C) Quantification of percent of cells containing autosomes with the indicated number of MLH1 foci in pachynema (same cells as presented in panel B).

### Sex chromosome pairing relies on ANKRD31–REC114 interactions

*Ankrd31*^*−/−*^ metaphase I cells frequently had unaligned chromosomes, and sex chromosomes failed to pair in 92% of otherwise normal looking pachytene cells (6, 7) (**Figure 6**). This high frequency of unpaired and/or achiasmate sex chromosomes is the likely cause of the highly penetrant metaphase I arrest in *Ankrd31*^*–/–*^ mutants.

**Figure 6.**
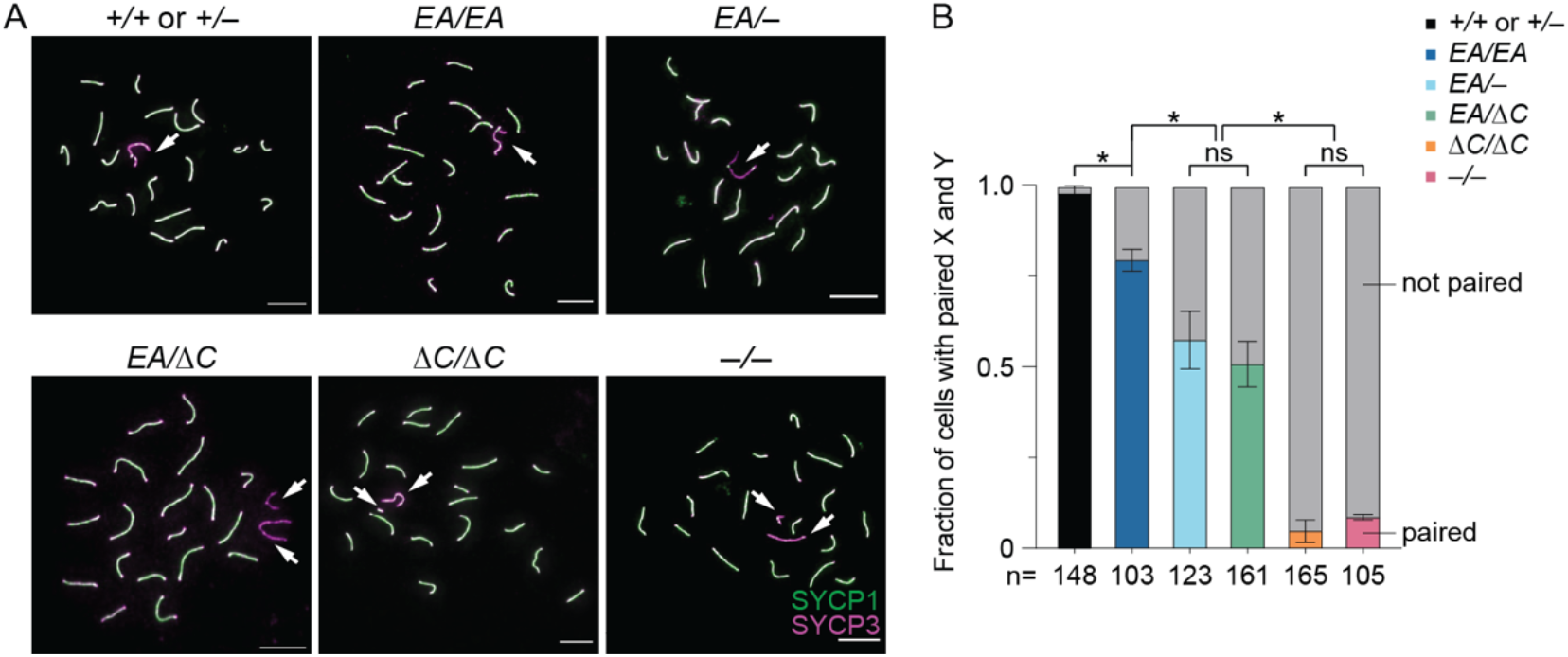
Sex chromosome pairing relies on ANKRD31–REC114 interactions. (A) Representative pachytene cell chromosome spreads stained with anti-SYCP3 and anti-SYCP1 antibodies. Scale bars, 10 μm. Arrowheads indicate sex chromosomes. (B) Quantification of X-Y pairing frequency at pachynema, based on immunofluorescence for SYCP3 and either SYCP1 or γH2AX, for three animals of each genotype. Total cell numbers are indicated under the graph. The results of Student’s t tests after arc sine transformation are shown: ns, not significant (p > 0.05), ∗p ≤ 0.05.

In keeping with the similarly high frequency of metaphase I arrest as *Ankrd31*^*−/−*^ (**Figure 2H**), *Ankrd31*^*ΔC/ΔC*^ spermatocytes showed 95.2% of sex chromosomes remaining unpaired in pachynema (**Figure 6**). In contrast, *Ankrd31*^*EA/EA*^ had a much more modest defect, with only 20.6% of sex chromosomes unpaired, while *Ankrd31*^*EA/–*^ and *Ankrd31*^*EA/ΔC*^ compound heterozygotes again had an intermediate defect, with about 50% of sex chromosomes unpaired (**Figure 6**). Therefore, the likelihood of successful sex chromosome pairing also correlates with the strength of the ANKRD31–REC114 interaction.

### Altered REC114 and ANKRD31 localization in interaction-deficient mice

ANKRD31 is required for the formation of immunostaining blobs of REC114 and other proteins on mo-2 minisatellite arrays on the PAR and elsewhere (Introduction). We therefore asked if the ANKRD31– REC114 interaction itself is specifically required. When chromosome spreads were stained for REC114, blobs were reduced in number and intensity in all of the *Ankrd31* mutants, with the severity of the defect correlating with the strength of interaction defect (**Figure 7A–C**). ANKRD31 blobs showed similar decreases (**Figure 7D–F**).

**Figure 7.**
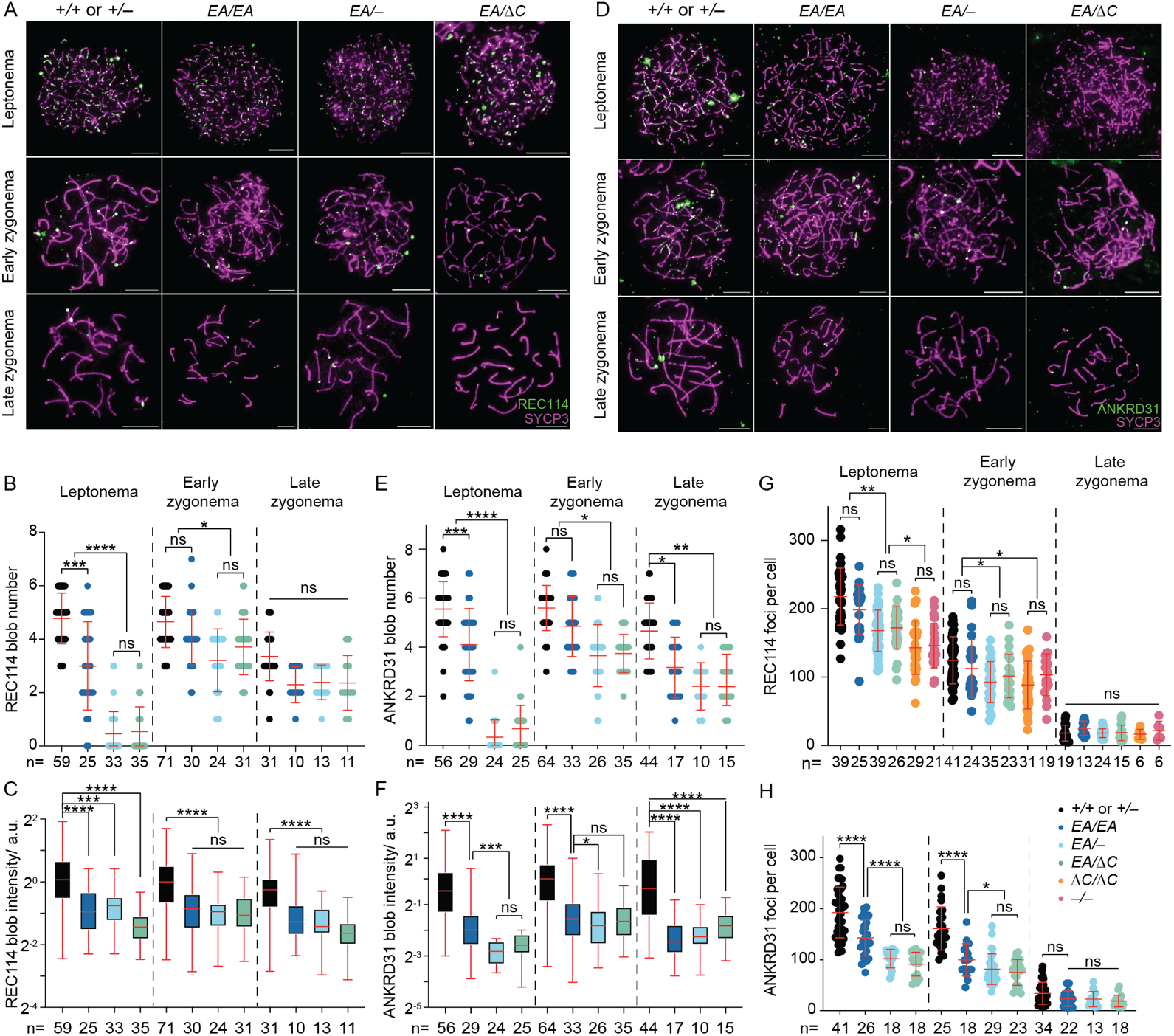
Altered REC114 and ANKRD31 localization in interaction-deficient mice. (A, D) Representative images of REC114 (A) or ANKRD31 staining of early prophase spermatocyte chromosome spreads. Scale bars, 10 μm. (B–C, E–F) Quantification of the REC114 blob number (B) and intensity (C); ANKRD31 blob number (E) and intensity (F); and numbers of small foci of REC114 (G) or ANKRD31 (H) from two or more animals of each genotype. The total cell numbers analyzed are given below each plot. Each point in the scatterplots is the measurement for one cell; red lines indicate means ± SD. In the boxplots (C, F), the intensities of blobs are normalized to the median blob intensity in the wild type in early zygonema. The boxes indicate the median, 25^th^, and 75^th^ percentiles. The whiskers indicate the 10^th^ and 90^th^ percentiles. The results of negative binomial regression tests (B, E) or two-tailed Mann-Whitney U tests (C, F, G, and H) are shown: ns, not significant (p > 0.05), ∗p ≤ 0.05, ∗∗p ≤ 0.01, ∗∗∗p ≤ 0.001, and ∗∗∗∗p ≤ 0.0001.

REC114 and ANKRD31 also extensively colocalize with one another in numerous small foci spread across the chromatin (6, 7). The small REC114 foci still form in the absence of *Ankrd31*, but are reduced in number and intensity during leptonema and early zygonema (6, 7) (**Figure 7A, G**). Not surprisingly then, the smaller REC114 foci were still present in our nonnull *Ankrd31* mutants. Fewer REC114 foci were observed in the truncation mutant at leptonema and early zygonema, similar to the pattern previously shown for *Ankrd31*^*−/−*^ spermatocytes. However, there was no obvious decrease in *Ankrd31*^*EA/EA*^ cells, and the compound heterozygotes again showed an intermediate phenotype, with a modest decrease in focus numbers (**Figure 7A, G**).

The small foci of ANKRD31 were more strongly affected by the *Ankrd31* mutations. *Ankrd31*^*EA/EA*^ cells had significantly fewer ANKRD31 foci at leptonema and early zygonema, and the compound heterozygotes were still further decreased (**Figure 7D, H**). Thus, even though the interaction-defective mutant proteins accumulate to normal levels in testes, their localization on chromatin is substantially altered. These findings confirm and extend prior observations and indicate that the ANKRD31–REC114 interaction is important for assembly both of the large immunostaining structures on mo-2 regions and of smaller foci genome-wide.

### The DSB landscape is shaped by the ANKRD31–REC114 interaction

In most mammals, DSB locations are controlled by PRDM9, which binds to specific DNA sequences and methylates nearby nucleosomes on histone H3 lysines 4 and 36, thereby targeting SPO11 (9). By singlestranded DNA sequencing (SSDS) of DMC1-bound DNA, most DSB hotspots genome wide overlap with sites of PRDM9-dependent H3K4me3 (10, 29). In the absence of *Prdm9*, DSBs are instead directed to other H3K4me3-modified loci (default hotspots) such as transcription promoters and CpG islands (29).

ANKRD31 is important for establishing normal DSB distributions (6, 7). *Ankrd31*^*−/−*^ spermatocytes have little or no SSDS signal in the PAR or PAR-proximal hotspots and there is a mixed usage of both PRDM9-targeted and default hotspots elsewhere in the genome (6, 7). Therefore, we asked if disruption of the ANKRD31–REC114 interaction would alter meiotic DSB locations.

To do this, we performed Exo7/T-seq (modified from S1-seq (30, 31) and END-seq (32)) to map DSBs genome-wide from testes of 14.5 dpp juvenile mice. In this method, high molecular weight genomic DNA embedded in agarose is digested with exonuclease VII and exonuclease T to remove the single-stranded DNA from resected DSBs, then sequencing adapters are ligated to the blunted DNA ends (**Figure S3A**). Two biological replicates were generated for each genotype and averaged **(Figure S3B)**.

In wild-type mice, Exo7/T-seq yielded the expected enrichment of top-strand reads to the right of PRDM9-specified hotspots and bottom strand reads to the left (reflecting 5’→3’ resection rightward and leftward, respectively) along with a central signal derived from recombination intermediates (**Figure S3A**) (31, 32). These resection and central signals were retained in *Ankrd31*^*−/−*^, consistent with previous SSDS experiments (6, 7), as well as in *Ankrd31*^*ΔC/ΔC*^ and the other *Ankrd31* mutant genotypes tested (**Figure 8A–B**). Interestingly, a modest (∼100 nucleotide) decrease in average resection lengths was observed in *Ankrd31*^*ΔC/ΔC*^ and *Ankrd31*^*−/−*^ (**Figures 8B–C and S3C–D**).

**Figure 8.**
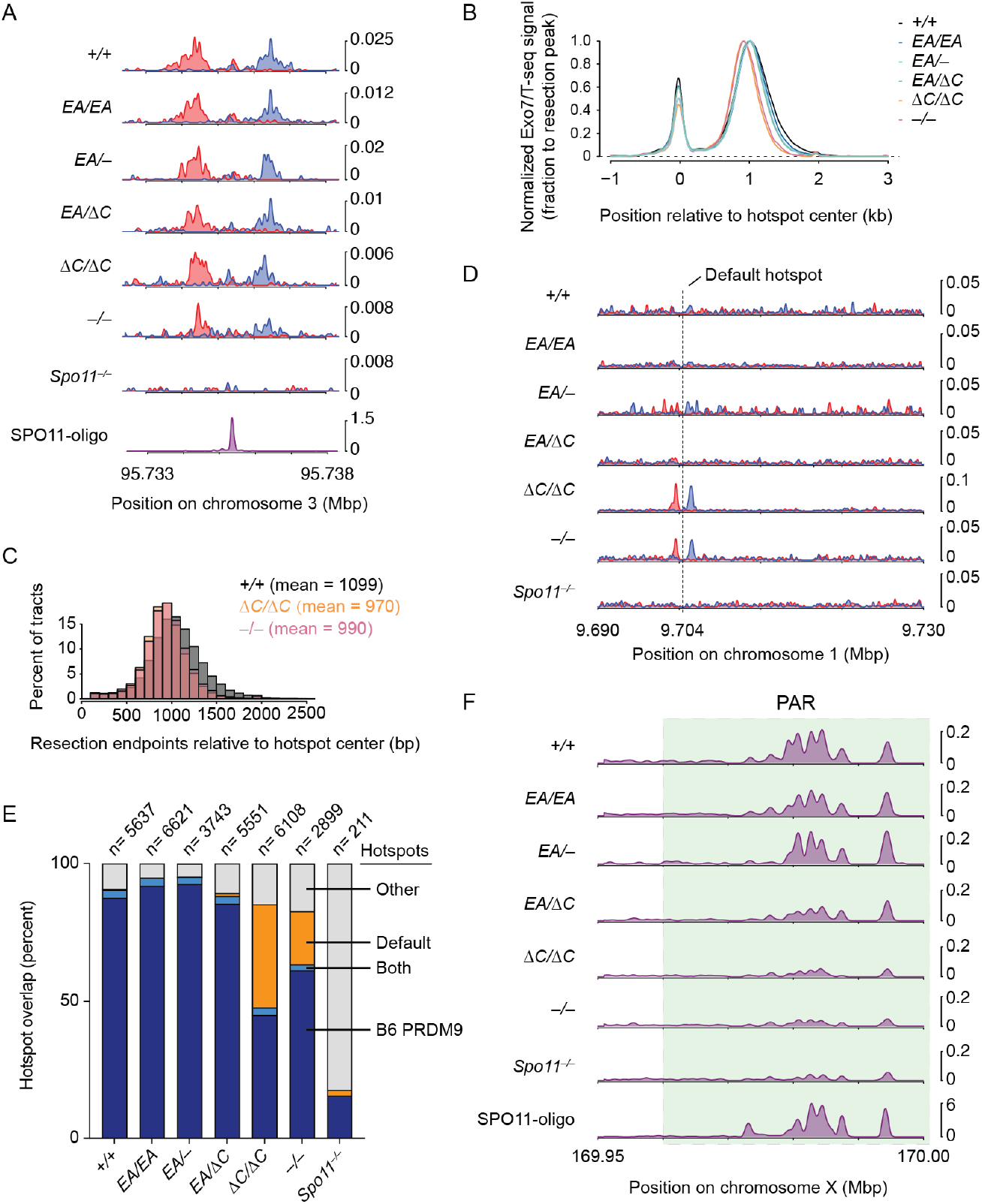
The DSB landscape is shaped by the ANKRD31–REC114 interaction. (A) Strand-specific Exo7/T-seq (reads per million mapped reads [RPM]) at a representative DSB hotspot. Top-strand reads are shown in blue, bottom-strand reads in red. Data are smoothed with a 151-bp Hanning window. The SPO11-oligo profile is shown below (33). (B) Averaged Exo7/T-seq signals around PRDM9-directed hotspots (n = 13,960 SPO11-oligo hotspots in the C57BL/6J strain (33)). The bottom-strand reads were flipped and combined with the top-strand reads and normalized to the peak height of resection endpoints (Methods and **Figure S3D**). (C) Comparison of resection length distributions. (D) Strand-specific Exo7/T-seq around a representative default hotspot. Data in 10 bp bins are smoothed with a 51-bin Hanning window. (E) Overlap of Exo7/T-seq peak calls in the indicated genotype with either PRDM9-directed SPO11-oligo hotspots (33) or the top 10,000 hottest default hotspots called using SSDS in *Prdm9*^*–/–*^ mice (10). Underlying data for peak calls and overlaps are provided in **Supplemental Table S1**. (F) Exo7/T-seq at the PAR (green shaded region). Sequencing reads from top and bottom strands were combined. Data in 40 bp bins are smoothed with a 51-bin Hanning window.

As previously shown using SSDS (6, 7), *Ankrd31*^*−/−*^ mutants displayed pronounced DSB signals at default hotspots in addition to PRDM9-directed hotspots (**Figure 8D**). *Ankrd31*^*ΔC/ΔC*^ likewise showed elevated use of default hotspots (**Figure 8D**), consistent with its close similarity to *Ankrd31*^*–/–*^ for other phenotypes.

To explore default hotspot usage more closely, we called peaks from Exo7/T-seq signals from each genotype indicated, and counted their overlap with either PRDM9-targeted hotspots (n=13,960 from SPO11 oligo sequencing (33)) or the 10,000 hottest of previously defined default hotspots (10) (**Figure 8E and Table S1**). As a control for meiotic DSB-independent background, we included *Spo11*^*–/–*^ mutants. *Ankrd31*^*−/−*^ and *Ankrd31*^*ΔC/ΔC*^ mutants showed an obvious elevation of default hotspot usage compared to other genotypes tested (**Figure 8E**). The *Ankrd31*^*EA/EA*^, *Ankrd31*^*EA/–*^ and *Ankrd31*^*EA/ΔC*^ mutants behaved broadly similarly to wild type.

*Ankrd31*^*−/−*^ mutants displayed little if any of the high-level Exo7/T-seq signals normally found in the PAR and were similar to the *Spo11*^*–/–*^ control (**Figure 8F**), consistent with prior SSDS studies (6, 7). *Ankrd31*^*ΔC/ΔC*^ mutants were similar (**Figure 8F**). This finding is consistent with the results that sex chromosomes rarely pair in these two mutants. In contrast, *Ankrd31*^*EA/EA*^ and the two compound heterozygotes displayed strong Exo7/T-seq signals in the PAR regions, broadly similar to control mice.

## Discussion

Although multiple axis-associated proteins are now known to promote or control meiotic DSB formation, recombination, and repair in mice, it remains poorly understood how the interactions among these proteins contribute to their functions. ANKRD31 emerged recently as playing a unique set of roles: on the PAR it is essential for high-level DSB formation, while globally it promotes normal timing and locations of DSBs but is dispensable for DSB formation per se. Because ANKRD31 interacts with multiple proteins that function in meiosis, including REC114, IHO1, MEI1, ZMYM3, and PTIP, it has been argued that it acts as a modular scaffold that anchors other proteins together and to meiotic chromosomes (6-8). Our findings are consistent with this interpretation and show that specifically disrupting the ANKRD31–REC114 interaction to different extents proportionately affected ANKRD31 function in vivo.

*Ankrd31*^*ΔC/ΔC*^ males showed phenotypes that were comparable to *Ankrd31*^*–/–*^ in every aspect that we characterized. It is striking that losing just the C-terminal 1.65% of the protein, with all of the tested interactions between ANKRD31 and its partners other than REC114 remaining intact, leads to a complete loss of function. This finding emphasizes the importance of the ANKRD31–REC114 interaction. Defects in *Ankrd31*^*EA/EA*^ mutants were comparatively much milder, even though the ANKRD31– REC114 interaction was severely disrupted as measured by Y2H assays. We speculate that multivalent interactions involving networks of ANKRD31-interacting proteins may compensate in vivo for attenuation (but not elimination) of the specific interaction with REC114.

ANKRD31 and REC114 are interdependent for their full level of chromatin association in both small foci and large blobs (6-8), and our results show that interactions of ANKRD31 with its other partners are not sufficient in the absence of REC114 interaction. This suggests that ANKRD31 and REC114 and their interaction are central to assembly of blobs on PAR and other mo-2 regions. In *Saccharomyces cerevisiae*, heterotrimeric Rec114–Mei4 complexes or homotetramers of Mer2 (homologous to IHO1 in mice) bind DNA in a highly cooperative manner in vitro, forming large nucleoprotein condensates containing many Rec114–Mei4 and mouse protein copies (34). A minimal trimerization domain from both yeast REC114–MEI4 is sufficient for condensate formation (35), suggesting a possible connection between the condensates characterized for the yeast proteins and the formation of small foci and blobs in vivo in mice. Although a detailed understanding of the mechanism of mo-2-associated blob formation remains elusive, we consider it likely that the multivalent ANKRD31 scaffold works interdependently with its binding partners to assemble condensates on mo-2 chromatin.

Since the function of ANKRD31 in the PAR is important in male but not female meiosis, ANKRD31 deficiency causes sexually dimorphic phenotypes: spermatogenesis fails catastrophically but oogenesis proceeds, albeit greatly diminished. In clinical studies, two heterozygous variants (p.Gln 329* and c.1565-2A>G (splice donor before exon 11)) in *ANKRD31* were found in patients with premature ovarian insufficiency (POI) (36). Both variants are predicted to result in early protein truncation that should disrupt ANKRD31 interactions with most or all of its known partners, including REC114. The POI resembles known features of the mouse *Ankrd31* null and truncation mutants, suggesting that ANKRD31 functions are conserved between mouse and human. Since these *ANKRD31* mutations cause POI when heterozygous, the truncated proteins may have dominant interfering effects on human oogenesis, or *ANKRD31* function may be dosage sensitive—analogous to but more pronounced than what we observed in mouse spermatogenesis.

Another woman has been reported to be homozygous for a C to T splicing donor mutation in *ANKRD31* (37). This mutation is predicted to cause a C-terminal truncation lacking exons 24 and 25 (106 amino acids), which is larger than our mouse *Ankrd31^ΔC^* truncation. No published clinical information is available for this individual, but our findings, combined with those of Wang et al. (2021), raise the possibility that she may be predisposed to POI. These clinical findings highlight the importance of understanding the molecular functions of ANKRD31 in meiosis.

## Materials and Methods

### Mice

Mice were maintained and euthanized under US federal regulatory standards. Experiments were approved by the Memorial Sloan Kettering Cancer Center (MSK) Institutional Animal Care and Use Committee (IACUC). Animals were fed regular rodent chow with ad libitum access to food and water. The previously described *Ankrd31*^*–*^ allele (*Ankrd31*^*em1Sky*^, MGI: 6343226) carries insertion of an A residue in exon 3, causing frameshift and premature termination (6). Interaction-deficient mice were generated by the MSKCC Mouse Genetics core facility by targeting exon 25. A guide RNA cassette with sequence (5’-GATACTTCAAATCAGCAAGC) and single-stranded donor DNA (5’-GAGTCGATGCAAACCATCCCTCATTATCTACAGATAAAAGAGATACTTCAAATCAGCAAGCAGGAGCTTCTGCCATGCCATGTAATGGAGCAGCATTGGAAGTTCTATGTGGGACGCTC) harboring desired missense mutations were microinjected using conventional techniques (38) into pronuclei of CBA/J × C57BL/6J F2 hybrid zygotes generated by crossing CBAB6F1/J hybrid females with C57BL/6J males. Founder mice (CBA/J × C57BL/6J F2 hybrid) were crossed to C57BL/J6 mice purchased from Jackson Laboratories to obtain germline transmission, then heterozygous animals were backcrossed to C57BL/J6 for at least 3 generations. The systematic names are *Ankrd31*^*em3Sky*^ for the EA allele (MGI: 7428356) and *Ankrd31*^*em4Sky*^ for the ΔC allele (MGI:7428358). ***Ankrd31EA/EA*** animals were generated by crossing ***Ankrd31+/EA*** heterozygous mice with ***Ankrd31+/EA*** heterozygous or ***Ankrd31EA/EA*** mice. ***Ankrd31***^*ΔC/ΔC*^ animals were generated by crossing ***Ankrd31+/****ΔC* heterozygous mice. ***Ankrd31–/–*** homozygous animals were generated as described (6). ***Mice were genotyped using*** PCR on cut toes lysed in ***Direct Tail lysis buffer (Viagen)*** with primers as indicated in **Supplemental Table S2**.

### Yeast two-hybrid assays

Genes were amplified by PCR from a testis cDNA library and digested by EcoR I (NEB) and Nde I (NEB), and then cloned into either pGADT7 or pGBKT7. Interaction deficient mutations were generated from corresponding vectors from ref. (6) containing wild-type sequences, using primers harboring desired mutations (**Supplemental Table S2**) (6). The ΔC construct was made by introducing the same 1-bp deletion as found in the mouse allele, so it fully mimics the truncation and extra amino acids encoded by frame shift. All plasmids were confirmed by sequencing of inserts. Mating of bait- and prey-containing strains (Y187 and Y2HGold yeast strains) and selection on synthetic dextrose plates lacking tryptophan and leucine and containing 200 ng/ml Aureobasidin A (SD-Trp/Leu/AbA) alone or also lacking adenine and histidine (SD-Trp/Leu/Ade/His/AbA) were performed following the manufacturer’s instructions (Clontech).

### Immunoprecipitation and immunoblot analysis of ANKRD31

Because of the reduced size and altered cellularity of testes from adult *Ankrd31*^*ΔC/ΔC*^ mice, we used testes of younger mutant mice (1.5 mos old) to assess protein levels (**Figure 1C, left**). We used adult testes for ANKRD31-EA (**Figure 1C, right**) because the missense mutation causes less of a reduction in testis weight. The slight decrease in ANKRD31-EA levels compared to the littermate control may be a consequence of the smaller testes in the mutant.

Decapsulated testes were snap-frozen in an Eppendorf tube in liquid nitrogen and stored at –80°C until usage. The frozen tissue was disrupted with a plastic pestle in RIPA buffer (50 mM Tris-HCl pH 7.5, 150 mM NaCl, 0.1% SDS, 0.5% sodium deoxycholate, 1% NP40) supplemented with protease inhibitors (Roche Mini tablets). The homogenate was supplemented with Benzonase nuclease (EMD Millipore (70664-3), 28 unit/ml) and 10 mM MgCl2 and incubated with end-over-end rotation for 1 hr at 4 °C. The samples were centrifuged at 21,130 g for 20 min at 4 °C. The supernatant was transferred to a new Lobinding tube (Eppendorf). The extract was pre-cleared with 50 µl protein A Dynabeads (Thermofisher) per sample by end-over-end rotation for 1 hour at 4 °C followed by magnetic removal of the beads. Then, 2 µg of antibodies (guinea pig anti-ANKRD31, rabbit anti-ANKRD31, or rabbit anti-REC114 (6)) were added to pre-cleared lysates and incubated with end-over-end rotation overnight at 4 °C. Protein A Dynabeads (50 µl) were added to the tubes and incubated for 1 hr with end-over-end rotation at 4 °C. Beads were washed three times with 500 µl RIPA buffer, eluted in 1× Nu-PAGE LDS sample buffer (Invitrogen) with 50 mM DTT, and incubated for 10 min at 70 °C.

Eluted proteins were separated on 3%–8% Tris-Acetate NuPAGE precast gels (Life Technologies) at 150 V for 70 min and were transferred to polyvinylidene difluoride (PVDF) membranes by wet transfer method in transfer buffer (192 mM glycine, 25 mM Tris, 10% methanol) at 120 V for 40 min at 4 °C. Membranes were blocked with 5% non-fat milk in 1× phosphate buffered saline (PBS) with 0.1% Tween (PBS-T) for 1 hr at room temperature. Blocked membranes were incubated with primary antibodies (guinea pig anti-ANKRD31, 1:4000; rabbit-ANKRD31, 1:4000) overnight at 4 °C. Membranes were washed with PBS-T for 3 × 10 min at room temperature on an orbital shaker, then incubated with HRP-conjugated secondary antibodies (rabbit anti-guinea pig IgG (Abcam 6771), 1:6000; goat anti-rabbit IgG (Biorad 170-6515)) for 1 hr at room temperature. Membranes were washed with PBS-T for 3 × 10 min and were developed by ECL Prime (GE Healthcare) and imaged on a Biorad ChemiDoc MP Imaging System.

### Spermatocyte chromosome spreads

Decapsulated testes were deposited in 50 ml Falcon tubes containing 2 ml TIM (104 mM NaCl, 45 mM KCl, 0.6 mM KH2PO4, 1.2 mM MgSO4, 6.0 mM sodium lactate, 1.0 mM sodium pyruvate, 0.1% glucose). Collagenase (200 μl of a 20 mg/ml solution in TIM) was added and incubated at 450 rpm for 55 min at 32 °C in a thermomixer. Samples were filled to a final volume of 15 ml with TIM, then centrifuged for 1 min at 39 g at room temperature. The supernatant was carefully removed with pipettes and the washing was repeated three times. Separated tubules were resuspended in 2 ml TIM, then 20 μl DNase I (400 μg/ml in TIM) and 200 μl trypsin (7 mg/ml in TIM) were added sequentially and incubated on a thermomixer for 15 min at 32 °C at 450 rpm. Trypsin inhibitor (500 μl of a 20 mg/ml solution in TIM) or FBS and 50 μl of the DNase I solution was added to terminate the reaction. A transfer pipet was used to further disperse the tissue by pipetting up and down for about 2 min. Cell suspensions were then filtered through a 70-μm cell strainer into a new 15 ml Falcon tube. TIM was added to a final volume of 15 ml and was centrifuged for 5 min at 108 g. The cell pellet was resuspended with 15 μl DNase I solution and TIM was added to a final volume of 15 ml. The washing procedure was repeated 2 times. The final pellet was resuspended in TIM according to the original testis weight (∼100 mg in 10 ml). This single-cell suspension (500 µl) was transferred to an Eppendorf tube and centrifuged for 3 min at 845 g. The cell pellet was resuspended in 40 µl of freshly prepared 0.1 M sucrose and incubated for 8 min at room temperature. Edges of superfrost glass slides were covered by Immedge pen, and each slide received 85 μl of 1% paraformaldehyde (PFA) (dissolved in presence of NaOH at 65 °C, 0.15% Triton, pH 9.3, filtered through a 0.22 μm filter, kept at –20 °C until usage). After incubation, 20 μl of cell suspension in the sucrose solution was added, slides were swirled three times, and dried in a closed slide box for 2 hr, followed by drying with a half-open lid for 1 hr at room temperature. Slides were washed in a Coplin jar on a shaker 1 × 5 min in milli-Q water and 3 × 5 min with 0.4% PhotoFlow (Kodak), then air-dried. Slides were stained immediately or were wrapped in aluminum foil and stored at −80°C.

### Immunostaining

Slides of spermatocyte chromosome spreads were blocked for 30 min at room temperature in a Coplin jar with 100 ml fresh block solution (1× PBS with 0.05% Tween-20 and 3 mg/ml bovine serum albumin (BSA)). Slides were incubated with 80 µl primary antibody in block solution and covered by parafilm overnight at 4 °C in a humid chamber. Slides were washed 3 × 5 min in block solution, then incubated with 80 µl secondary antibody in block solution for 30 min in a humid chamber at room temperature. Slides were then washed 3 × 5 min in the dark with block solution, rinsed in milliQ H2O, and mounted with Vectashield containing DAPI. Primary and secondary antibodies are listed in **Supplemental Table S3**.

### Histology

Testes and epididymides dissected from adult mice were fixed in Bouin’s fixative for 4 to 5 hr at room temperature, or in 4% PFA overnight at 4 °C. Bouin’s fixed testes were washed in 15 ml milliQ H2O on a horizontal shaker for 1 hr at room temperature, followed by five 1-hr washes in 15 ml of 70% ethanol on a roller at 4 °C. PFA-fixed tissues were washed 4 × 5 minutes in 15 ml milliQ H2O at room temperature. Fixed tissues were stored in 70% ethanol before embedding in paraffin and sectioning (5 μm for testes, 8 μm for ovaries). The tissue sections were deparaffinized with EZPrep buffer (Ventana Medical Systems). Hematoxylin and eosin (H&E) staining, periodic acid Schiff (PAS) staining and immunohistochemical TUNEL assay were performed by the MSK Molecular Cytology Core Facility using the Autostainer XL (Leica Microsystems, Wetzlar, Germany) automated stainer for H&E with hematoxylin counterstain, and using the Discovery XT processor (Ventana Medical Systems, Oro Valley, Arizona) for TUNEL. The detection was performed with DAB detection kit (Ventana Medical Systems) according to manufacturer’s instructions. Slides were counterstained with hematoxylin and coverslips were mounted with Permount (Fisher Scientific).

### Image acquisition

Images of spread spermatocytes were acquired on a Zeiss Axio Observer Z1 Marianas Workstation, equipped with an ORCA-Flash 4.0 camera, illuminated by an X-Cite 120 PC-Q light source, with 100× 1.4 NA oil immersion objective. Marianas Slidebook (Intelligent Imaging Innovations, Denver Colorado) software was used for acquisition.

Whole slides (histology), either H&E or TUNEL stained, were scanned and digitized with the Panoramic Flash Slide Scanner (3DHistech, Budapest, Hungary) with a 20× 0.8 NA objective (Carl Zeiss, Jena, Germany). High-resolution images of H&E and IHC images were acquired with a Zeiss Axio Imager microscope using a 63× 1.4 NA oil immersion objective (Carl Zeiss, Jena, Germany).

### Image analysis

To quantify foci of ANKRD31, REC114, DMC1, RAD51, RPA, and MLH1, single cells were manually cropped and counted in Fiji. For ANKRD31 and REC114 blobs, overlap with SYCP3 and fluorescence intensity were determined with a thresholding algorithm regardless of meiotic stage, and a difference of Gaussian (DoG) blur was used to isolate REC114 or ANKRD31 foci. Scripts are available on Github: https://github.com/Boekhout/ImageJScripts. All images were manually inspected upon analysis.

For γH2AX intensity analysis, cells were sub-staged manually by SYCP3 staining. Regions of interest were manually drawn and integrated density was measured. A region containing no cells was measured for background subtraction. To combine measurements from different experiments, the integrated density was normalized to the mean integrated intensity of wild-type leptotene cells within each experiment.

### DNA extraction in agarose plugs

Testes from 14.5-dpp juvenile mice were decapsulated and incubated in DMEM containing 0.1% polyvinyl alcohol (PVA, Sigma), 0.1% BSA (Gibco) with 1 mg/ml collagenase type IV (Worthington) and 1 mg/ml Dispase II (Sigma) for 20 min at 35 °C in a thermomixer at 450 rpm. Seminiferous tubules were then rinsed three times and further treated with trypsin (TrypLE™ express enzyme, Gibco) and 1 μg/ml DNase I (Roche) for 15 min at 35 °C in a thermomixer at 450 rpm. Trypsin was inactivated with 5% FBS and tubules were further dissociated by gentle pipetting. Cells were passed through a 70-μm cell strainer (BD Falcon) and washed three times in GBSS containing 0.1% PVA.

Cells were embedded in plugs of 1% low-melting-point agarose (Lonza) in GBSS (1.5 million to 2 million cells per plug). Plugs were incubated with 100 μg/ml proteinase K (Roche) in lysis buffer (0.5 M EDTA at pH 8.0, 1% N-lauroylsarcosine sodium salt) at 50 °C over two nights. Plugs were washed 5 × 20 min with TE (10 mM Tris-HCl at pH 7.5, 1 mM EDTA at pH 8.0), and then incubated with 100 μg/ml RNase A (Thermo) for 3 hr at 37 °C. Plugs were then washed five times with TE and stored in TE at 4 °C until usage.

### Exo7/T-seq

Exo7/T-seq combines elements of previously described S1-seq (30, 31) and END-seq (32) methods. Specifically, it uses *E. coli* exonuclease VII and exonuclease T to remove ssDNA, like END-seq, but uses the methods for genomic DNA preparation, adapter ligation, and sequencing library preparation from S1-seq. Agarose plugs containing genomic DNA prepared as previously described (31) were washed in 1 ml of Exo VII buffer (50 mM Tris-HCl, 50 mM sodium phosphate, 8 mM EDTA, 10 mM 2-mercaptoethanol, pH 8.0) 2 × 15 min, equilibrated with 50 U of exonuclease VII (NEB) in 100 μl of Exo VII buffer for 10 min on ice and then incubated for 60 min at 37 °C using a thermomixer at 400 rpm. Plugs were rinsed with TE and then washed with 1 ml NEB buffer 4 (NEB) 3 × 15min, equilibrated with 75 U of exonuclease T (NEB) on ice for 30 min, and then incubated for 90 min at 24 °C using a thermomixer at 400 rpm. Plugs were washed in 500 µl of 1× T4 polymerase buffer (1× T4 ligase buffer (NEB) supplemented with 100 μg/ml BSA and 100 μM dNTPs (Roche)) 4 × 30 min on ice. The wash solution was then replaced with 1× T4 polymerase buffer containing 30 U T4 DNA polymerase (NEB) and incubated at 12 °C for 30 min. P5 adapters were ligated to the ends with 1 μl of 2000 U/μl T4 DNA Ligase (NEB) at 16 °C for 20 hr. After ligation, plugs were washed in 1 ml TE three times and incubated in TE overnight at 4 °C.

Plugs were washed with 500 μl of 1× β-agarase I buffer (10 mM Bis-Tris-HCl, 1 mM EDTA, pH 6.5) on ice for 30 min, then 150 μl of 1× β-agarase I buffer was added and incubated at 70 °C for 5 min to melt the agarose, followed by vortexing for 30 s and brief centrifugation. The vortex and spin-down procedure was repeated three times. Samples were cooled to 42 °C and 2 μl of β-agarase I enzyme (NEB) was added and incubated at 42 °C for 90 min with occasional vortexing. DNA was sheared into fragment sizes ranging 200–500 bp with a Covaris system (E220 Focused-ultrasonicator, microtube-500) using the following parameters: delay 300 s then three cycles of [peak power 175, duty factor 20, cycles/burst 200, duration 30 s, and delay 90 s]. Sonicated samples were added to 1250 μl of 100% ethanol and 55 μl of 3 M sodium acetate and incubated overnight at –20°C.

Ethanol precipitated DNA was dissolved in 52 μl of TE for 1 hr at 37 °C. SPRIselect beads (Beckman Coulter) were used to remove unligated adapters. Fragments containing the biotinylated adapter were purified with Dynabeads™ M-280 streptavidin (Thermo Fisher). The end-repair reaction was done using the End-it DNA end-repair kit (Lucigen). P7 adapters were ligated to DNA fragments as described above. PCR was done on beads. PCR products were purified with 0.9× AMPure XP beads (Beckman Coulter) to remove primer dimers and unligated adapters. DNA was sequenced on the Illumina HiSeq platform in the Integrated Genomics Operation at MSK. We obtained paired-end reads of 50 bp. Bioinformatic analysis was performed as described previously (30, 31). In brief, Trim Galore <http://www.bioinformatics.babraham.ac.uk/projects/trim_ga-lore/> was used to trim and filter reads with the arguments -paired -length 15. Bowtie2 (39) was run with the arguments -N 1 -X 1000 to map sequence reads onto the mouse reference genome (mm10). Picard <**Error! Hyperlink reference not valid**.> eliminated reads that were duplicated. With the -q 20 argument, Samtools (40) extracted properly mapped and uniquely mapped reads (MAPQ ≥ 20). Reads for Exo7/T-seq were counted at the nucleotide immediately adjacent to the position where the biotinylated adapter DNA was mapped (i.e., corresponding to the last position of the ssDNA). Maps were analyzed using R (versions 3.3.1 and 4.0.3). Exo7/T-seq signal from the top and bottom strands was averaged and co-oriented and plotted in R for each genotype. The averaged profile was smoothed with a 151-bp Hanning filter. Differences in background levels between samples, likely because of variation in testis cellularity, cause genotype-independent variation in absolute signal level (31) (**Figure S3D**). Therefore, to compare spatial patterns of resection in **Figure 8B**, the average profile for each genotype was normalized to the peak height of resection endpoints. To do this, an estimated background was removed by subtracting the value 2,500 bp away from the hotspot center. The background-subtracted profile was then normalized to the maximum value between 100 and 2,500 bp. Negative values were set as zero for plotting purposes.

To plot the histogram of resection lengths, an estimated background value was removed by subtracting the signal 2.5 kb away from the hotspot center. By setting values for positions 100 bp and >2.5 kb to zero, the signal near and farther from the hotspot core was eliminated. Fractions of remaining total signal were calculated every 100 bp and plotted.

Uniquely mapping fragments derived from Exo7/T-seq were used to identify hotspot locations (peak calling). Peak calling was performed using MACS (v.2.2.7.1) (41) with the following parameters: -g mm -- keep-dup all – nolambda -- nomodel. Numbers of peak calls are indicated in **Supplemental Table S1**. Narrow peaks were then widened by 5 kb and merged. Then merged peaks were compared with PRDM9-targeted hotspots (n=13,960 from SPO11 oligo sequencing) or the 10,000 hottest of previously defined default hotspots (10, 33) to calculate the fraction of hotspot usage. Overlaps were defined as peaks with a shared region of at least 1 bp.

### Data availability

Exo7T-seq sequencing data have been deposited in the Gene Expression Omnibus (GEO) repository under accession number GSE229450. Underlying data for all plots, including exact p values, are provided in **Supplemental File 1**.

## Acknowledgments

This article is subject to the Open Access to Publications policy of the Howard Hughes Medical Institute (HHMI). HHMI lab heads have previously granted a nonexclusive CC BY 4.0 license to the public and a sublicensable license to HHMI in their research articles. Pursuant to those licenses, the author-accepted manuscript of this article can be made freely available under a CC BY 4.0 license immediately upon publication.

We thank D. Ontoso, M. Marcet, M. Arter, and K. Liu of the Keeney laboratory for discussions and experimental advice. We thank the MSK Molecular Cytology core facility (N. Fan and M. Pulina) for histology; the Integrated Genomics Operation (IGO) for sequencing; and the Mouse Genetics core facility (P. Romanienko and W. Mark) for generating the *Ankrd31* mutant mouse lines. MSK core facilities are supported by National Cancer Institute Cancer Center support grant P30 CA08748. The IGO was further funded by the Cycle for Survival and the Marie-Josée and Henry R. Kravis Center for Molecular Oncology. This work was supported by a Basic Research Innovation award from MSK (to S.K. and D. Patel) and NIH grants R35 GM118092 (to S.K.), and R01 HD110120 (to S.K. and D. Patel).

## Notes

### Competing Interest Statement

The authors have declared no competing interest.

### Summary of Updates

Added background information about female meiosis; added information about biological replicates for deep sequencing experiments and changed S1-seq normalization procedure; text changes throughout to improve clarity and correct typos; supplemental files updated.

