## Supplemental Figures and Tables for "Essential roles of the ANKRD31-REC114 interaction in meiotic recombination and mouse spermatogenesis"

**This pdf contains:**

Supplemental Figures S1-S3
Supplemental Tables S1-S3

**Additional information, in separate .xls file:**
Supplemental File 1

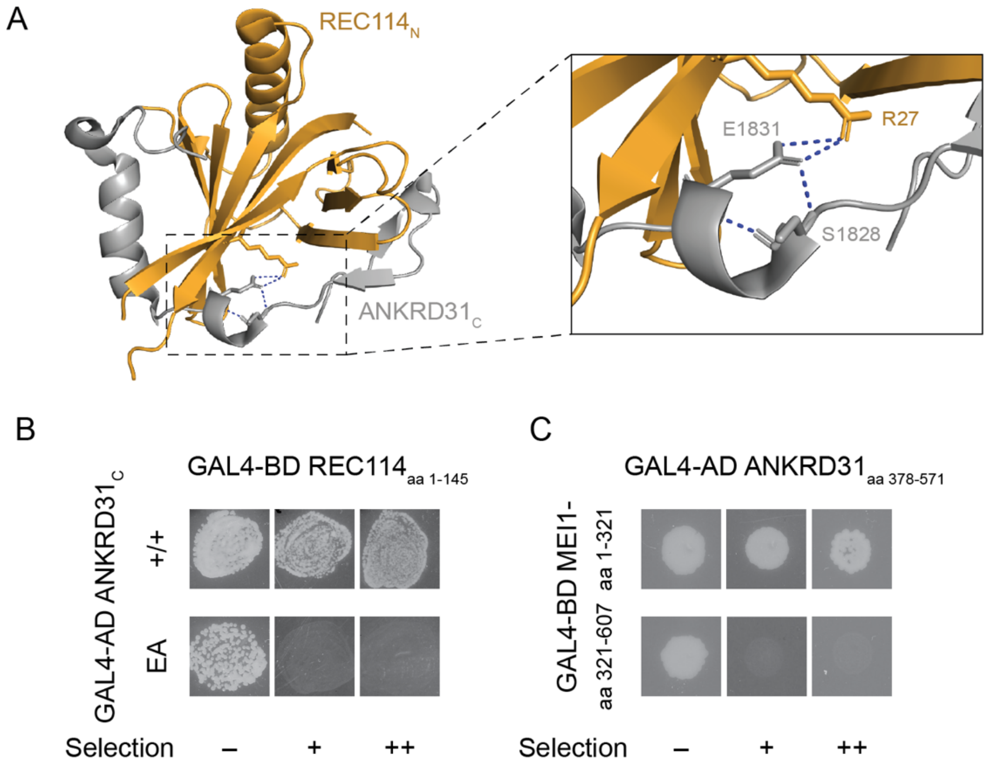


**Figure S1. ANKRD31 interactions with REC114 and MEI1.** (A) Crystal structure of ANKRD31$_C$ (gray) bound to the REC114$_N$ pleckstrin homology domain (yellow) (PDB: 6NXF) (Boekhout et al., 2019). The zoomed detail highlights contacts made by ANKRD31 Glu-1831. (B) Y2H interactions of REC114$_{aa\ 1\text{-}158}$ and wild type or EA mutant ANKRD31$_{C.}$ (C) Y2H interactions of full-length and fragments of ANKRD31 and MEI1. Cells express the indicated Gal4 activating domain (AD) and binding domain (BD) fusions. EV, empty vector. "Selection" indicates amino acid dropouts and aureobasidin to detect reporter activation at moderate (+) and high (++) stringency.

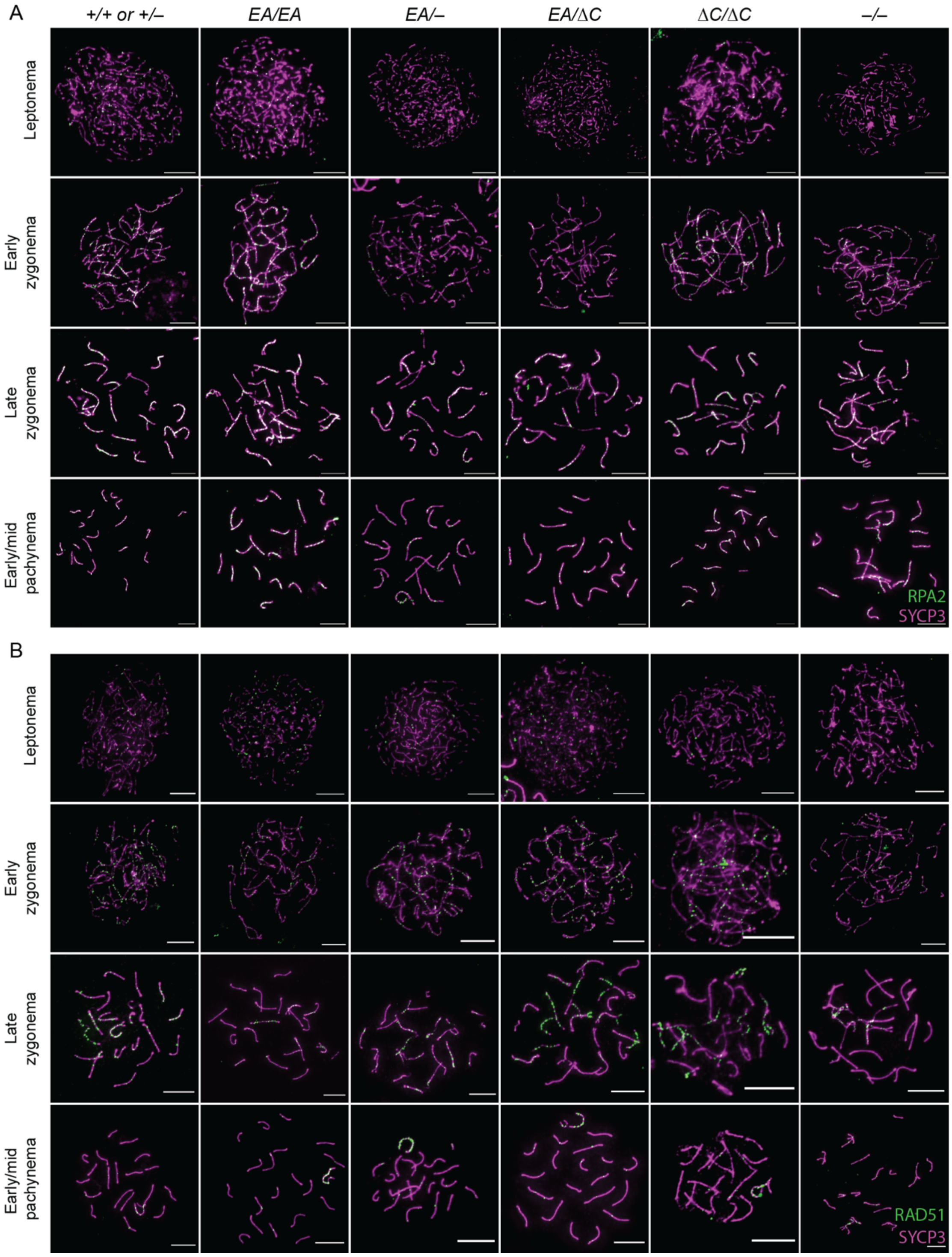


**Figure S2. ANKRD31–REC114 interaction deficiencies cause progressive defects in DSB formation and recombination.** Representative images of RPA2 staining (A) or RAD51 staining (B) of spermatocyte chromosome spreads. Quantification is presented in **Figure 7**. Scale bars, 10 μm.

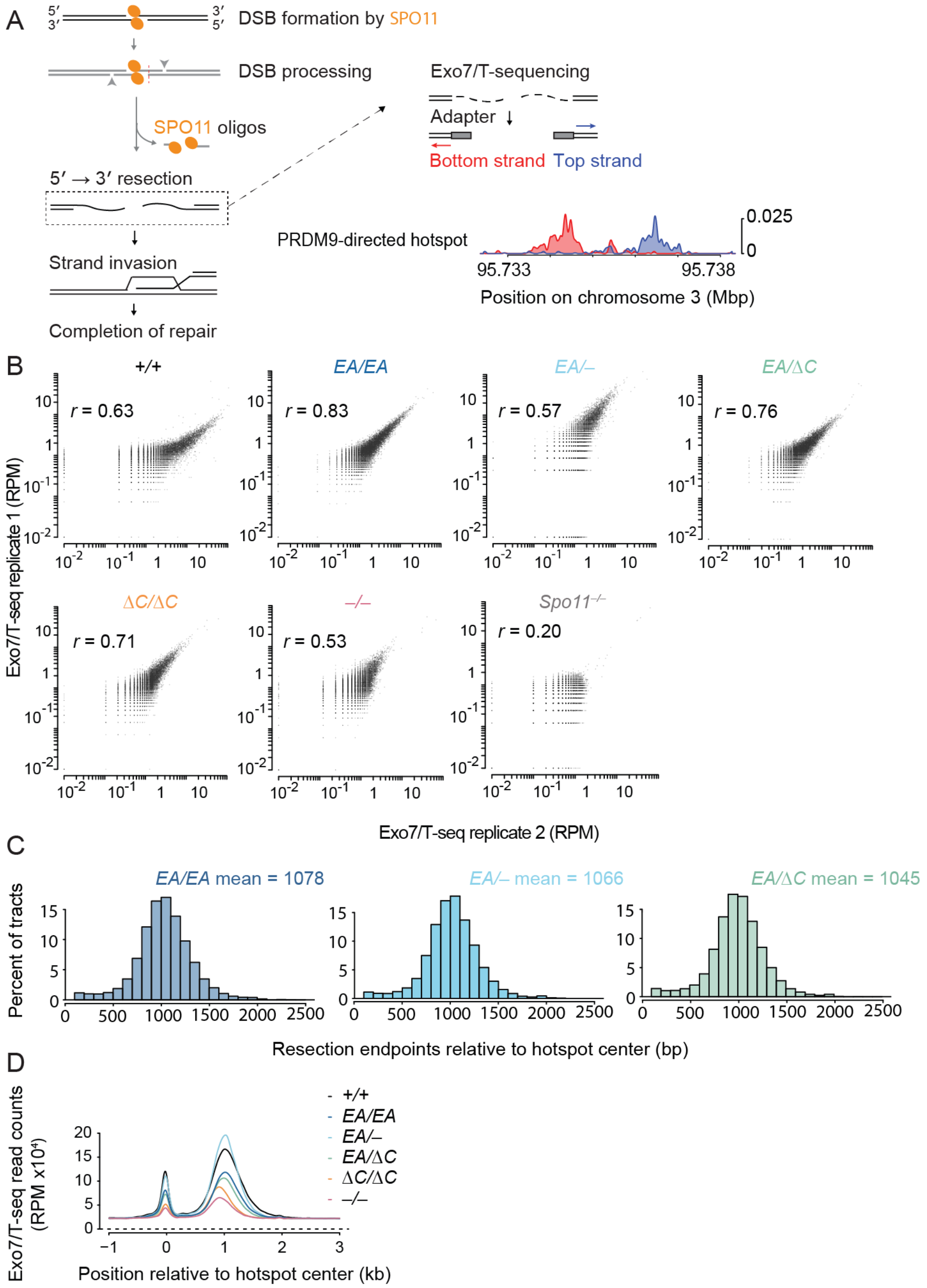


**Figure S3. Schematic of Exo7/T sequencing and resection length**. (A) Steps in DSB formation and processing and schematic of Exo7/T sequencing to detect resection endpoints. An example profile is shown from wild type at a PRDM9-directed hotspot in B6 mice. (B) Correlation (Pearson's *r*) of Exo7/T-seq read counts between biological replicates. Each point is a SPO11-oligo hotspot, with Exo7/T-seq signal summed from −2000 to −250 bp (bottom strand) and +250 to +2000 bp (top strand) around hotspot centers. Hotspots with ≤ 0 RPM were excluded for Pearson's *r* calculation. Hotspots with ≤ $10^{-2}$ RPM were set as $10^{-2}$ for plotting purposes only. Note that the correlation for the *Spo11*$^{-/-}$ replicates is poor because the sequencing signal represents mostly dispersed, nonspecific background. (C) Resection length distributions in

*Ankrd31*$^{EA/EA}$, *Ankrd31*$^{EA/–}$ and *Ankrd31*$^{EA/\Delta C}$. (D) Non-normalized, averaged Exo7/T-seq signals around PRDM9-directed hotspots (n = 13,960 SPO11-oligo hotspots in the C57BL/6J strain (33)). The bottom-strand reads were flipped and combined with the top-strand reads.

**Supplemental Table S1. Overlap of Exo7/T-seq peak calls with SPO11-directed or default hotspots.**

**Overlap of Exo7/T-seq peak calls with SPO11-directed or default hotspots**

| | Total peak called | Merged peaks | Overlap with SPO11-directed hotspots | Overlap with default hotspots | Overlap with both | Overlap with neither |
|---|---|---|---|---|---|---|
| *Ankrd31*$^{+/+}$ | 5637 | 2704 | 2443 | 86 | 77 | 252 |
| *Ankrd31*$^{EA/EA}$ | 6621 | 3213 | 3046 | 98 | 96 | 165 |
| *Ankrd31*$^{EA/–}$ | 3743 | 2075 | 1975 | 59 | 57 | 98 |
| *Ankrd31*$^{EA/\Delta C}$ | 5551 | 2931 | 2583 | 111 | 80 | 317 |
| *Ankrd31*$^{\Delta C/\Delta C}$ | 6108 | 3562 | 1700 | 1432 | 99 | 529 |
| *Ankrd31*$^{–/–}$ | 2899 | 1872 | 1187 | 400 | 41 | 326 |
| *Spo11*$^{–/–}$ | 211 | 97 | 15 | 2 | 0 | 80 |

**Supplemental Table S2. Primers for genotyping and cloning.**

| **Primers for genotyping** | | |
|---|---|---|
| Genotype | Primer Name | Primer Sequence |
| *Ankrd31*$^{EA}$ | 18MGAnkF08 | 5'-CCGATCACGTATGCTCTGATACGG -3' |
| | 18MGAnkR05 | 5'-ACTGACCATACGAGTTATAGTGCAGG -3' |
| *Ankrd31*$^{\Delta C}$ | 18MGAnkF05 | 5'-AGCAGGTACTTCCAGAGAGTCGAT -3' |
| | 18MGAnkR03 | 5'-GCTAACTAAGCTACCAAGAAAGCAGAGC -3' |

| **pGAD-ANKRD31 cloning primers** | | |
|---|---|---|
| *Ankrd31*$^{EA}$ | | 5'-CTTCAAATCAGCAAGCAGGCGC-3' |
| | | 5'-CATTACATGGCATGGCAGAAGCGCC-3' |
| *Ankrd31*$^{\Delta C}$ | | 5'-CAGAAGCAGGAGCTTCTGCCATGCCATGTAATGGAGCAG-3' |
| | | 5'-AGAAGCTCCTGCTTCTGATTTGAAGTATCTCTTTTATCTGT-3' |

**Supplemental Table S3. Antibodies.**

**Primary Antibodies**

| Target | Dilution for IF | Dilution for immunoblot | Host species | Supplier | Catalog number |
|---|---|---|---|---|---|
| γH2AX | 1:6000 | n/a | Rabbit | Abcam | ab2893 |
| DMC1 (H100) | 1:100 | n/a | Rabbit | Santa Cruz | sc-22768 |
| MLH1 | 1:25 | n/a | Mouse | BD-Pharmingen | 51-1327GR |
| RPA2 | 1:1000 | n/a | Rabbit | Abcam | ab76420 |
| RAD51 | 1:100 | n/a | Rabbit | Aviva | ARP33450 |
| SYCP3 (D-1) | 1:300 | n/a | Mouse | Santa Cruz | sc-74569 |
| SYCP3 | 1:200 | n/a | Rabbit | Nova | NB300-232 |
| SYCP1 | 1:200 | n/a | Rabbit | Abcam | ab15090 |
| REC114 | 1:200 | n/a | Rabbit | Stanzione et al., 2016 | n/a |
| ANKRD31 | 1:200 | 1:4000 | Guinea Pig | Boekhout et al., 2019 | n/a |

**Secondary Antibodies**

| Target | Dilution for IF | Fluorophore | Host species | Supplier | Catalog number |
|---|---|---|---|---|---|
| anti Guinea pig | 1:400 | Alexa-488 | Goat | Mol Probes | A-11073 |
| anti Mouse | 1:400 | Alexa-594 | Goat | Mol Probes | A-11005 |
| anti Mouse | 1:400 | Alexa-594 | Donkey | Life Technologies | A-21203 |
| anti Mouse | 1:400 | Alexa-488 | Donkey | Life Technologies | A-21202 |
| anti Rabbit | 1:400 | Alexa-594 | Goat | Invitrogen | A-11012 |
| anti Rabbit | 1:400 | Alexa-488 | Goat | Invitrogen | A11034 |
| anti Rabbit | 1:400 | Alexa-488 | Donkey | Thermofisher Scientific | R37118 |
